# Repurposing the Antiviral Agent Pibrentasvir: *In Vitro* Synergistic Effects in Combination with Different Azole Antifungal Agents

**DOI:** 10.64898/2026.07.30.741933

**Authors:** Sijie Liu, Jing Hu, Min Shen, Lu Ge, Hanmo Yang, Xinyi Tao, Heng Zhang, Yi Sun

**Affiliations:** Department of Dermatology, Hubei Provincial Clinical Research Center for Diagnosis and Therapeutics of Pathogenic Fungal Infection, Jingzhou Hospital Affliated to Yangtze University, Jingzhou, Hubei Province, 434100, China; Department of Clinical Medicine, Xiangyang Polytechnic University, Xiangyang, Hubei Province, 441050, China; Department of Otolaryngology, Hubei Provincial Clinical Research Center for Diagnosis and Therapeutics of Pathogenic Fungal Infection.Jingzhou Hospital Affiliated to Yangtze University, Jingzhou, Hubei Province, 434100, China; Department of Clinical Medicine, Yangtze University, Jingzhou, Hubei Province, 434100, China

**Keywords:** pibrentasvir, azoles, *in vitro*, synergistic effect

## Abstract

**Objective:** To investigate the combined effects of multiple drugs and provide more therapeutic options for invasive fungal infections, this study evaluated the *in vitro* susceptibility of *Aspergillus* spp., *Candida auris*, *Cryptococcus neoformans*, and *Exophiala dermatitidis* to pibrentasvir (PIB) in combination with itraconazole (ITR), voriconazole (VOR), posaconazole (POS), or fluconazole (FLU).

**Methods:** According to the M27-A3 and M38-A2 guidelines established by the Clinical and Laboratory Standards Institute (CLSI), the *in vitro* antifungal activities of PIB combined with ITR, VOR, POS, or FLU against 78 clinical isolates, including *Aspergillus* spp., *E. dermatitidis*, *C. auris*, and *C. neoformans*, were determined. The minimum inhibitory concentrations (MICs) and fractional inhibitory concentration indices (FICIs) were calculated to evaluate the synergistic effects.

**Results:** PIB alone exhibited no antifungal activity. Significant synergistic effects were observed when PIB was combined with azole antifungal agents. The PIB-POS combination showed synergistic effects against *Aspergillus* spp. (27/41, 65.90%), *C. auris* (9/10, 90.00%), *C. neoformans* (2/9, 22.20%), and *E. dermatitidis* (11/18, 61.11%). The PIB-ITR combination also showed synergistic effects against *Aspergillus* spp. (18/41, 43.9%), *C. auris* (9/10, 90.0%), *C. neoformans* (2/9, 22.2%), and *E. dermatitidis* (10/18, 55.5%). Synergistic effects were less frequently observed with the PIB-VOR or PIB-FLU combinations, and no antagonistic effects were observed.

**Conclusion:** This study demonstrates that PIB acts as an azole sensitizer, with the strongest synergistic effects observed when combined with POS or ITR, providing a new research direction for combination therapy against invasive fungal infections.

## Background

As the number of immunocompromised individuals continues to increase, together with the more frequent use of broad-spectrum antibiotics and immunosuppressive agents, both the incidence and mortality burden of invasive fungal infections (IFIs) have risen, making IFIs one of the major challenges in global public health. The high mortality associated with IFIs is comparable to that of common bacterial and parasitic diseases, such as tuberculosis and malaria. Approximately 90% of deaths attributable to fungal infections are caused by species of *Aspergillus* spp., *Candida*, or *Cryptococcus*^1^. In addition, infections caused by *Exophiala dermatitidis* should not be overlooked.

Airborne fungi of the genus *Aspergillus* can cause severe invasive infections in humans and are associated with substantial mortality, which may reach 90% in immunocompromised patients^2^. *Aspergillus fumigatus* is the most common causative species, accounting for approximately 90% of human infections^3^. It is estimated to cause up to 16 million pulmonary infections annually, resulting in hundreds of thousands of deaths^2,4^. However, *A. fumigatus* is not the only pathogenic species within this genus; *Aspergillus terreus*, *Aspergillus flavus*, and other species can also cause fatal disease. Since the first case of *Candida auris* was reported in Japan in 2009, infections caused by this pathogen have been documented in more than 40 countries, with reported mortality rates ranging from 30% to 60%. *C. auris* has become an important global nosocomial pathogen. Unlike other *Candida* species, it exhibits a unique tropism for the skin and can persistently colonize this site, thereby facilitating nosocomial persistence and clonal transmission and contributing to large-scale hospital outbreaks^5^. Both *Cryptococcus neoformans* and *E. dermatitidis* can cause fatal infections of the central nervous system. *C. neoformans* is widely distributed worldwide in association with pigeon droppings, and cryptococcal meningoencephalitis caused by this organism is one of the most common fungal infections of the central nervous system^6^. *E. dermatitidis* is widely distributed in warm and humid environments, including dishwashers, bathrooms, saunas, moist soil, and decaying plant material. Central nervous system infection caused by this organism represents one of its most severe and frequently fatal clinical manifestations^7^.

First-line azole antifungal agents inhibit sterol 14α-demethylase (CYP51), thereby blocking ergosterol biosynthesis, disrupting fungal cell membrane integrity, and inhibiting fungal growth^8^. However, antifungal resistance continues to emerge and spread, posing an important and increasingly serious threat to human health^9^. Consequently, combination antifungal therapy has become a key strategy aimed at achieving pharmacodynamic and pharmacokinetic synergy, shortening treatment duration, reducing drug dosages, broadening the antifungal spectrum, and delaying the emergence of resistance^10^.

Pibrentasvir (PIB) is one of the first-line agents used for the treatment of hepatitis C virus (HCV) and exerts its antiviral activity by inhibiting the NS5A protein involved in viral replication. Velpatasvir (VEL), which has a similar mechanism of action, has been shown to markedly enhance the antifungal activity of amphotericin B (AmB), demonstrating favorable synergistic effects. Moreover, the combination of VEL and AmB did not exacerbate AmB-associated cytotoxicity^11^. We therefore hypothesized that PIB could act as a sensitizer to azole antifungal agents.

To further investigate the potential of PIB-azole combinations and provide a new strategy for antifungal therapy, the present study evaluated the *in vitro* synergistic antifungal effects of PIB in combination with itraconazole (ITR), posaconazole (POS), voriconazole (VOR), or fluconazole (FLU) against 78 clinical isolates of *Aspergillus* spp., *C. auris*, *E. dermatitidis*, and *C. neoformans*, in accordance with the M27-A3 and M38-A2 guidelines issued by the Clinical and Laboratory Standards Institute (CLSI).

## Materials and Methods

### Fungal Strains, Antifungal Agents, and Chemical Reagents

A total of 78 fungal isolates were included in this study, all of which were obtained from the Hubei Provincial Clinical Research Center for the Diagnosis and Treatment of Pathogenic Fungal Infections. These included 41 *Aspergillus* isolates, comprising 27 *A. fumigatus* isolates (AF2-8 and AF16-35), five *A. flavus* isolates, five *A. terreus* isolates, and four *Aspergillus niger* isolates; 10 *C. auris* isolates (AR381-390), originally obtained from the CDC and FDA Antibiotic Resistance Isolate Bank; nine *C. neoformans* isolates; and 18 *E. dermatitidis* isolates.

All isolates were identified by morphological characterization and sequencing of the ITS and D1/D2 regions. The sequences were deposited in GenBank under accession numbers PP069948-PP070390. Before the experiments, all isolates were inoculated onto Martin agar medium and cultured at 35°C for 2-3 days for activation^12^. *Candida parapsilosis* ATCC 22019 and *A. flavus* ATCC 204304 were used as quality-control strains for antifungal susceptibility testing. PIB (CAS No. 1353900-92-1; purity ≥ 98%), POS (CAS No. 171228-49-2; purity > 99%), VOR; (CAS No. 137234-62-9; purity ≥ 98%), ITR (CAS No. 84625-61-6; purity > 98%), and FLU (CAS No. 86386-73-4; purity > 98%) were purchased from Aladdin. All compounds were dissolved in dimethyl sulfoxide (DMSO).

### *In Vitro* Combination Susceptibility Testing

The experimental procedures were performed in accordance with CLSI M27-A3^13^ and M38-A2^14^ guidelines. Fungal suspensions were prepared in sterile 0.9% sodium chloride solution. The conidial concentrations of *Aspergillus* spp. and *E. dermatitidis* were adjusted to 2-5 × 10^6^ conidia/mL, whereas the cell concentrations of *C. auris* and *C. neoformans* were adjusted to 2-5 × 10^5^ cells/mL^15^. The detailed experimental procedures were carried out according to the method described by Jia et al^16^. Briefly, the procedure was performed as follows. In a 96-well microtiter plate, 50 μL of serially diluted antifungal agents was dispensed along the columns. The final concentration ranges were 0.063-8 μg/mL for VOR and ITR, 0.031-4 μg/mL for POS, and 0.063-1 μg/mL for FLU. Subsequently, 50 μL of serially diluted PIB was dispensed across the rows, with a final concentration range of 0.125-8 μg/mL. Thereafter, 100 μL of fungal suspension was added to each well, resulting in final inoculum concentrations of 2-5 × 10^6^ conidia/mL for *Aspergillus* spp. and *E. dermatitidis*, and 2-5 × 10^5^ cells/mL for *C. auris* and *C. neoformans*. The plates were incubated at 35°C for 24 h for *C. auris* and *C. neoformans*, 48 h for *Aspergillus* spp., and 72 h for *E. dermatitidis*. For *Aspergillus* spp. and *E. dermatitidis*, the minimum inhibitory concentration (MIC) was defined as the lowest drug concentration that completely inhibited visible fungal growth^15^. For *C. auris* and *C. neoformans*, the MIC was defined as the lowest drug concentration that resulted in ≥ 50% inhibition of growth^12^.

MIC endpoints were determined by visual assessment of fungal growth, and the MIC values of each drug tested alone and in combination were recorded. The fractional inhibitory concentration index (FICI) was calculated using the following formula: FICI = (MIC of drug A in combination/MIC of drug A alone) + (MIC of drug B in combination/MIC of drug B alone). Drug interactions were interpreted as synergistic when the FICI was ≤ 0.50, indifferent when the FICI was > 0.50 and ≤ 4.00, and antagonistic when the FICI was > 4.00^17,18^. All experiments were independently repeated three times on separate days.

## Result

### *In Vitro* Interactions of PIB in Combination with Azoles against *Aspergillus* spp

PIB alone exhibited no detectable antifungal activity against any of the 41 *Aspergillus* isolates tested, with MICs of > 16 μg/mL. The MIC ranges of ITR, VOR, and POS were 0.125-8 μg/mL, 0.25-4 μg/mL, and 0.063-2 μg/mL, respectively. Synergistic interactions were observed for the PIB-ITR, PIB-VOR, and PIB-POS combinations against 18 (43.9%), 2 (4.9%), and 27 (65.9%) isolates, respectively. No antagonistic interactions were detected for any of the combinations (Table 1).

**Table 1.** *In vitro* drug sensitization results of PIB combined with azoles against *Aspergillus spp*.

| Strains | Alone |  |  |  | Combination |  |  |
| --- | --- | --- | --- | --- | --- | --- | --- |
|  | PIB | ITR | VOR | POS | PIB/ITR | PIB/VOR | PIB/POS |
| *A. fumigatus* |  |  |  |  |  |  |  |
| AF2 | >16 | 1 | 1 | 0.25 | 2/0.25(S) | 2/0.25(S) | 1/0.063(S) |
| AF3 | >16 | 1 | 0.25 | 0.5 | 2/0.25(S) | 0.125/0.25(I) | 0.5/0.125(S) |
| AF4 | >16 | 1 | 0.25 | 0.5 | 2/0.25(S) | 0.125/0.25(I) | 1/0.125(S) |
| AF5 | >16 | 1 | 0.25 | 0.25 | 4/0.25(S) | 0.125/0.25(I) | 2/0.063(S) |
| AF6 | >16 | 1 | 1 | 0.25 | 4/0.25(S) | 2/0.5(I) | 2/0.063(S) |
| AF7 | >16 | 1 | 0.5 | 0.5 | 2/0.25(S) | 0.25/0.25(I) | 1/0.125(S) |
| AF8 | >16 | 0.125 | 4 | 2 | 0.125/0.125(I) | 4/2(I) | 0.5/1(I) |
| AF16 | >16 | 0.5 | 0.25 | 0.25 | 4/0.125(S) | 8/0.125(I) | 2/0.063(S) |
| AF17 | >16 | 1 | 0.25 | 0.25 | 2/0.25(S) | 0.125/0.25(I) | 2/0.0625(S) |
| AF18 | >16 | 1 | 1 | 0.5 | 8/0.25(S) | 0.5/0.5(I) | 2/0.25(I) |
| AF19 | >16 | 1 | 1 | 0.25 | 2/0.25(S) | 1/0.25(S) | 2/0.031(S) |
| AF20 | >16 | 1 | 1 | 0.5 | 2/0.5(I) | 0.25/1(I) | 2/0.125(S) |
| AF21 | >16 | 1 | 1 | 0.25 | 4/0.25(S) | 4/0.5(I) | 2/0.063(S) |
| AF22 | >16 | 1 | 0.5 | 0.25 | 8/8(I) | 0.125/0.5(I) | 1/0.063(S) |
| AF23 | >16 | 8 | 4 | 0.5 | 4/0.25(S) | 0.125/4(I) | 1/0.125(S) |
| AF24 | >16 | 1 | 0.25 | 0.25 | 2/0.5(I) | 8/0.125(I) | 1/0.063(S) |
| AF25 | >16 | 1 | 0.5 | 0.5 | 4/0.25(S) | 0.125/0.5(I) | 2/0.125(S) |
| AF26 | >16 | 1 | 0.25 | 0.25 | 2/0.25(S) | 0.125/0.25(I) | 1/0.063(S) |
| AF27 | >16 | 1 | 0.5 | 0.25 | 2/0.25(S) | 4/0.25(I) | 2/0.063(S) |
| AF28 | >16 | 1 | 0.25 | 0.125 | 2/0.25(S) | 0.125/0.25(I) | 4/0.031(S) |
| AF29 | >16 | 8 | 2 | 0.063 | 8/8(I) | 0.5/1(I) | 0.125/0.063(I) |
| AF30 | >16 | 1 | 0.25 | 0.25 | 2/0.25(S) | 0.125/0.25(I) | 0.25/0.063(S) |
| AF31 | >16 | 1 | 0.25 | 0.125 | 2/0.25(S) | 0.125/0.25(I) | 2/0.031(S) |
| AF32 | >16 | 1 | 0.25 | 0.25 | 1/0.5(I) | 0.125/0.25(I) | 1/0.063(S) |
| AF33 | >16 | 1 | 0.25 | 0.125 | 1/0.5(I) | 2/0.125(I) | 1/0.031(S) |
| AF34 | >16 | 1 | 0.25 | 0.25 | 2/0.5(I) | 0.125/0.25(I) | 1/0.063(S) |
| AF35 | >16 | 1 | 0.25 | 0.125 | 2/0.25(I) | 0.125/0.25(I) | 0.5/0.063(S) |
| *A. flavus* |  |  |  |  |  |  |  |
| AFL2 | >16 | 0.5 | 0.5 | 0.25 | 0.125/0.5(I) | 0.5/1(I) | 4/0.75(I) |
| AFL13 | >16 | 1 | 1 | 0.25 | 0.5/0.5(I) | 0.125/1(I) | 2/0.125(I) |
| AFL22 | >16 | 1 | 1 | 0.5 | 1/1(I) | 0.125/1(I) | 1/0.25(I) |
| AFL23 | >16 | 0.5 | 1 | 0.25 | 1/0.5(I) | 0.125/1(I) | 2/0.125(I) |
| AFL24 | >16 | 1 | 2 | 0.5 | 0.125/0.5(I) | 0.125/2(I) | 0.5/0.25(I) |
| *A. terreus* |  |  |  |  |  |  |  |
| 453 | >16 | 0.5 | 1 | 0.5 | 0.125/0.5(I) | 0.5/2(I) | 0.5/0.125(S) |
| 455 | >16 | 0.25 | 0.5 | 0.125 | 4/0.5(I) | 0.125/0.5(I) | 1/0.063(I) |
| 465 | >16 | 0.25 | 1 | 0.25 | 8/0.5(I) | 1/0.5(I) | 1/0.063(S) |
| 479 | >16 | 0.25 | 0.5 | 0.125 | 0.125/0.5(I) | 0.125/0.5(I) | 0.5/0.063(I) |
| 503 | >16 | 0.25 | 1 | 0.25 | 0.5/0.5(I) | 2/0.5(I) | 4/0.063(S) |
| *A. niger* |  |  |  |  |  |  |  |
| AN2 | >16 | 1 | 1 | 0.25 | 0.125/1(I) | 2/0.5(I) | 2/0.125(I) |
| AN3 | >16 | 1 | 1 | 0.25 | 0.125/1(I) | 2/0.5(I) | 2/0.125(I) |
| AN4 | >16 | 2 | 1 | 0.25 | 1/4(I) | 0.125/1(I) | 0.125/0.25(I) |
| AN5 | >16 | 1 | 1 | 0.5 | 2/0.5(I) | 1/0.5(I) | 0.5/0.25(I) |
| Quality control |  |  |  |  |  |  |  |
| ATCC204304 | >16 | 1 | 1 | 0.25 | 0.125/1(I) | 0.125/1(I) | 1/0.125(I) |
| ATCC22019 | >16 | 0.25 | 0.125 | 0.063 | 0.5/0.063(S) | 0.125/0.125(I) | 1/0.031(I) |

Note: ITR, itraconazole; VOR, voriconazole; POS, posaconazole; PIB, pibrentasvir; S, synergy (FICI $\leq 0.5$ ); I, indifference (no interaction, FICI from $>0.5$ to $\leq 4$ ). MICs were the concentrations that achieved 100% growth inhibition; FICI: fractional inhibitory concentration index.

Further analysis revealed distinct patterns of synergy among the PIB-azole combinations. PIB-POS exhibited the broadest and strongest synergistic activity, particularly against *A. fumigatus* (24/27, 88.9%) and *A. terreus* (3/5, 60.0%). PIB-ITR showed synergistic activity against 18 of 27 *A. fumigatus* isolates (66.7%), whereas PIB-VOR was synergistic against only 2 of 27 isolates (7.4%). These findings indicate that PIB-POS exhibited more extensive synergistic activity than PIB-ITR and PIB-VOR across the tested *Aspergillus* species, with no detectable antagonism (Figure 1).

**Figure 1.**
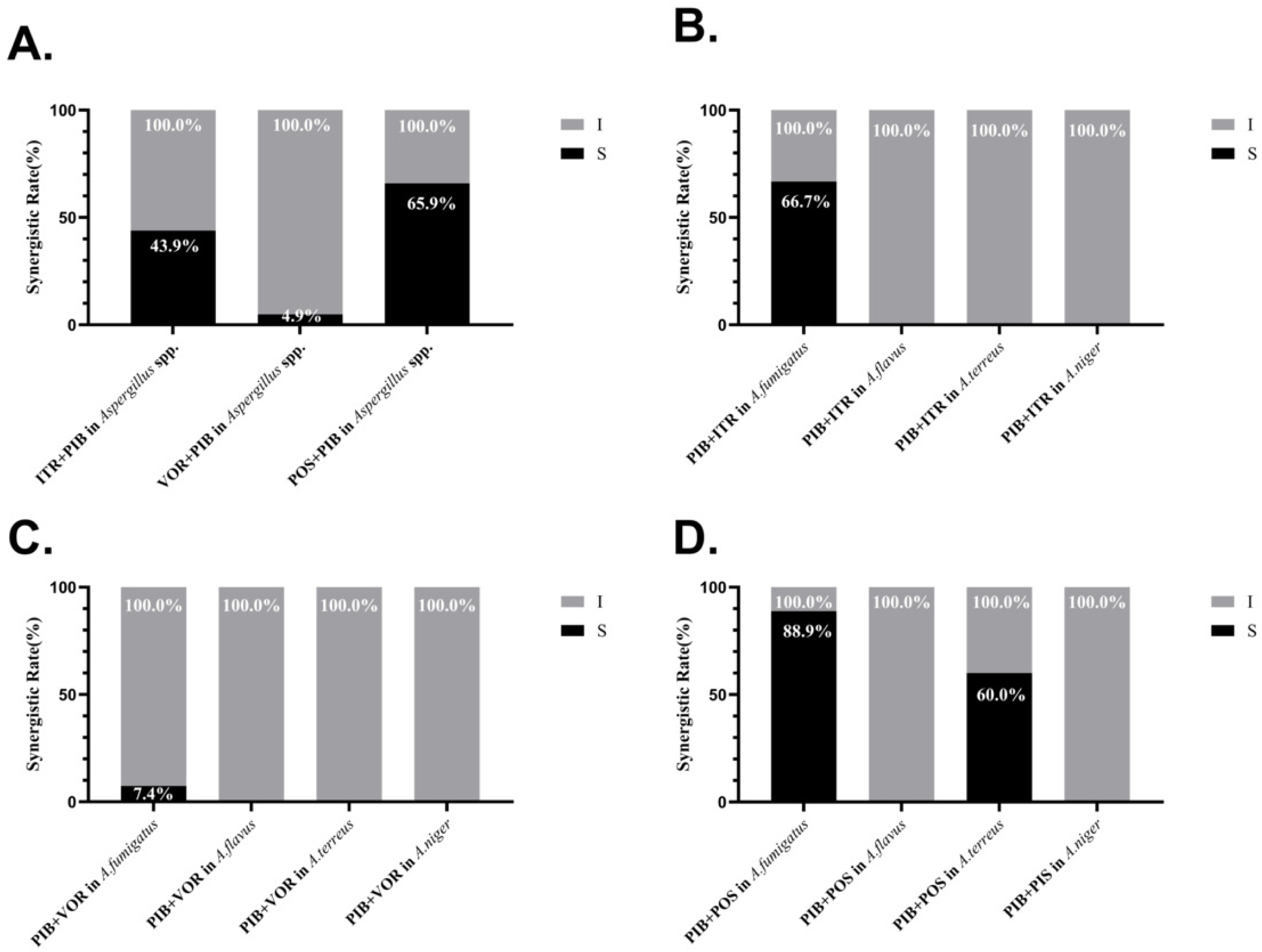
*In vitro* drug sensitization results of PIB combined with azoles against *Aspergillus* spp. Summary of drug interaction for the combination of PIB and azoles. (A) Summary of interaction relationships for all drug combinations against *Aspergillus* spp. (B-D) The fraction of in vitro interaction results of PIB combined with total result in *Aspergillus,* ITR, VOR, and POS antifungal agents, respectively.S, synergy (FICI of < 0.5); I, no interaction (indifference)(0.5 < FICI < 4).

### *In Vitro* Interactions of PIB in Combination with Azoles against *C. auris*

PIB alone showed no detectable antifungal activity against any of the 10 *C. auris* isolates, with MICs of > 16 μg/mL. The MIC ranges of ITR, VOR, and POS were 0.25-1 μg/mL, 0.125-1 μg/mL, and 0.125-0.25 μg/mL, respectively (Table 2). Synergistic interactions were observed for PIB-ITR, PIB-VOR, and PIB-POS against 9 (90.0%), 6 (60.0%), and 9 (90.0%) isolates, respectively. No antagonistic interactions were detected between PIB and any of the three azoles. These results demonstrate that PIB in combination with triazoles exhibited substantial synergistic activity against *C. auris*, particularly when combined with ITR or POS (Figure 2).

**Figure 2.**
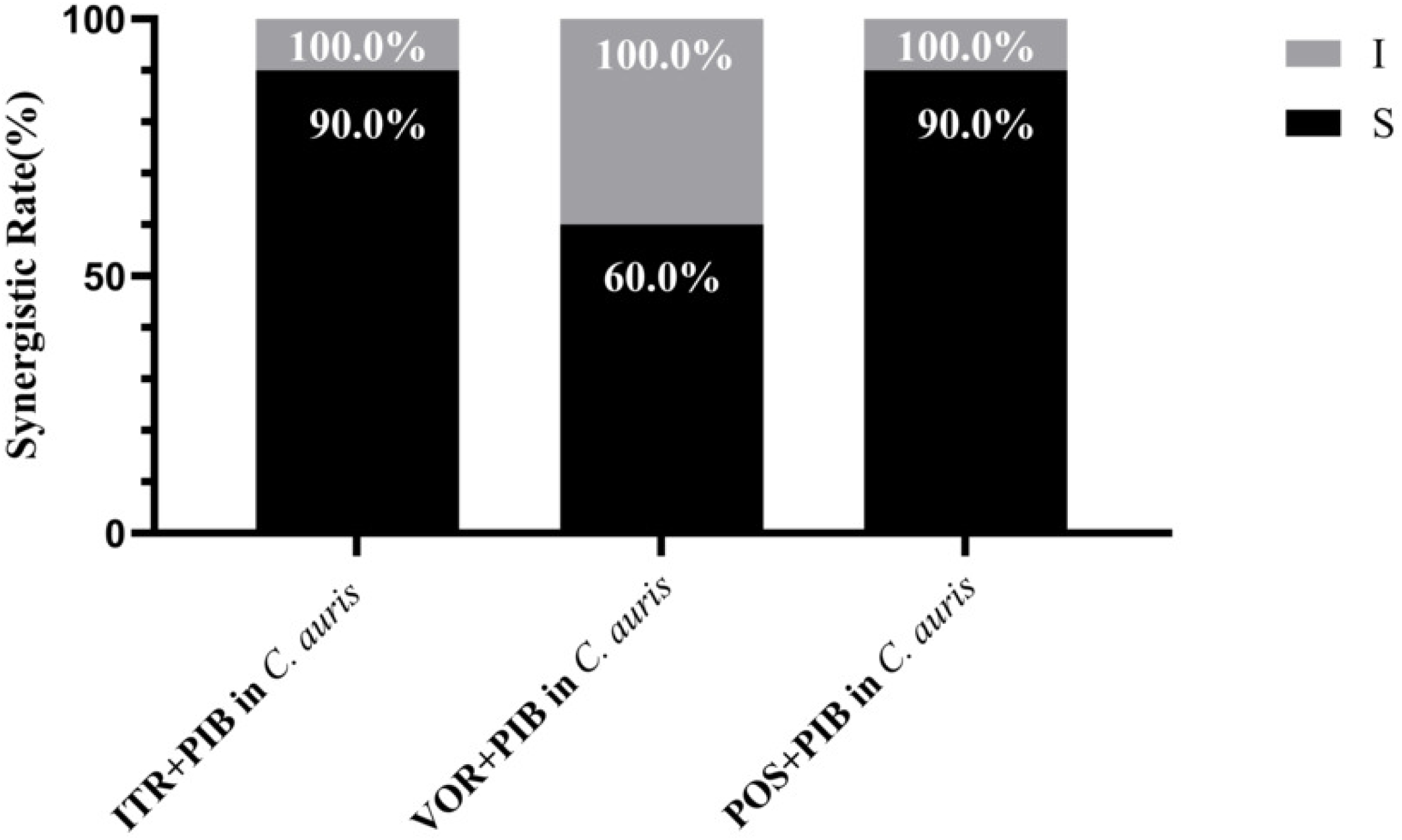
*In vitro* drug sensitization results of PIB combined with azoles against *C.aruis.* Summary of drug interaction for the combination of PIB and azoles. Summary of interaction relationships for all drug combinations against *C.aruis.* The fraction of in vitro interaction results of PIB combined with total result in *C.aruis,* ITR, VOR, and POS antifungal agents, respectively.S, synergy (FICI of < 0.5); I, no interaction (indifference)(0.5 < FICI < 4).

**Table 2.** *In vitro* drug sensitization results of PIB combined with azoles against *C. aruis*.

| Strains | Alone |  |  |  | Combination |  |  |
| --- | --- | --- | --- | --- | --- | --- | --- |
|  | PIB | ITR | VOR | POS | PIB/ITR | PIB/VOR | PIB/POS |
| AR381 | >16 | 0.25 | 0.125 | 0.125 | 1/0.063(S) | 0.25/0.063(I) | 1/0.031(S) |
| AR382 | >16 | 0.25 | 0.5 | 0.125 | 2/0.063(S) | 2/0.125(S) | 1/0.031(S) |
| AR383 | >16 | 0.25 | 0.5 | 0.125 | 1/0.063(S) | 1/0.125(S) | 2/0.031(S) |
| AR384 | >16 | 0.25 | 0.25 | 0.125 | 1/0.063(S) | 2/0.063(S) | 4/0.031(S) |
| AR385 | >16 | 0.5 | 0.5 | 0.25 | 2/0.125(S) | 0.5/0.25(I) | 2/0.031(S) |
| AR386 | >16 | 0.5 | 1 | 0.125 | 0.5/0.25(I) | 2/0.25(S) | 1/0.031(S) |
| AR387 | >16 | 1 | 0.5 | 0.25 | 0.5/0.25(S) | 4/0.125(S) | 0.5/0.125(I) |
| AR388 | >16 | 0.5 | 0.5 | 0.125 | 4/0.125(S) | 0.25/1(I) | 1/0.031(S) |
| AR389 | >16 | 0.25 | 0.125 | 0.125 | 2/0.063(S) | 0.5/0.063(I) | 2/0.031(S) |
| AR390 | >16 | 0.25 | 0.25 | 0.125 | 2/0.063(S) | 2/0.063(S) | 1/0.031(S) |
| **Quality control** |  |  |  |  |  |  |  |
| ATCC22019 | >16 | 0.25 | 0.125 | 0.063 | 0.5/0.063(S) | 0.125/0.125(I) | 1/0.031(I) |
| ATCC204304 | >16 | 0.5 | 0.5 | 0.25 | 0.125/0.5(I) | 0.125/0.5(I) | 0.5/0.125(I) |

Note: ITR, itraconazole; VOR, voriconazole; POS, posaconazole; PIB, pibrentasvir; S, synergy (FICI $\leq 0.5$ ); I, indifference (no interaction, FICI from $>0.5$ to $\leq 4$ ). MICs were the concentrations that achieved 100% growth inhibition; FICI: fractional inhibitory concentration index.

### *In Vitro* Interactions of PIB in Combination with Azoles against *C. neoformans*

PIB alone exhibited no detectable antifungal activity against any of the nine *C. neoformans* isolates, with MICs of > 16 μg/mL. The MIC ranges of ITR, VOR, POS, and FLU were 0.125-0.5 μg/mL, 0.125 μg/mL, 0.125-0.25 μg/mL, and 0.25-1 μg/mL, respectively. Synergistic interactions were observed for PIB-ITR, PIB-POS, and PIB-FLU against 3 (33.3%), 2 (22.2%), and 4 (44.4%) isolates, respectively, whereas no synergy was observed for PIB-VOR. No antagonistic interactions were detected for any of the combinations (Table 3). These findings indicate that PIB combined with azole antifungal agents exhibited synergistic activity against a subset of *C. neoformans* isolates, with PIB-FLU showing the highest rate of synergy (Figure 3).

**Figure 3.**
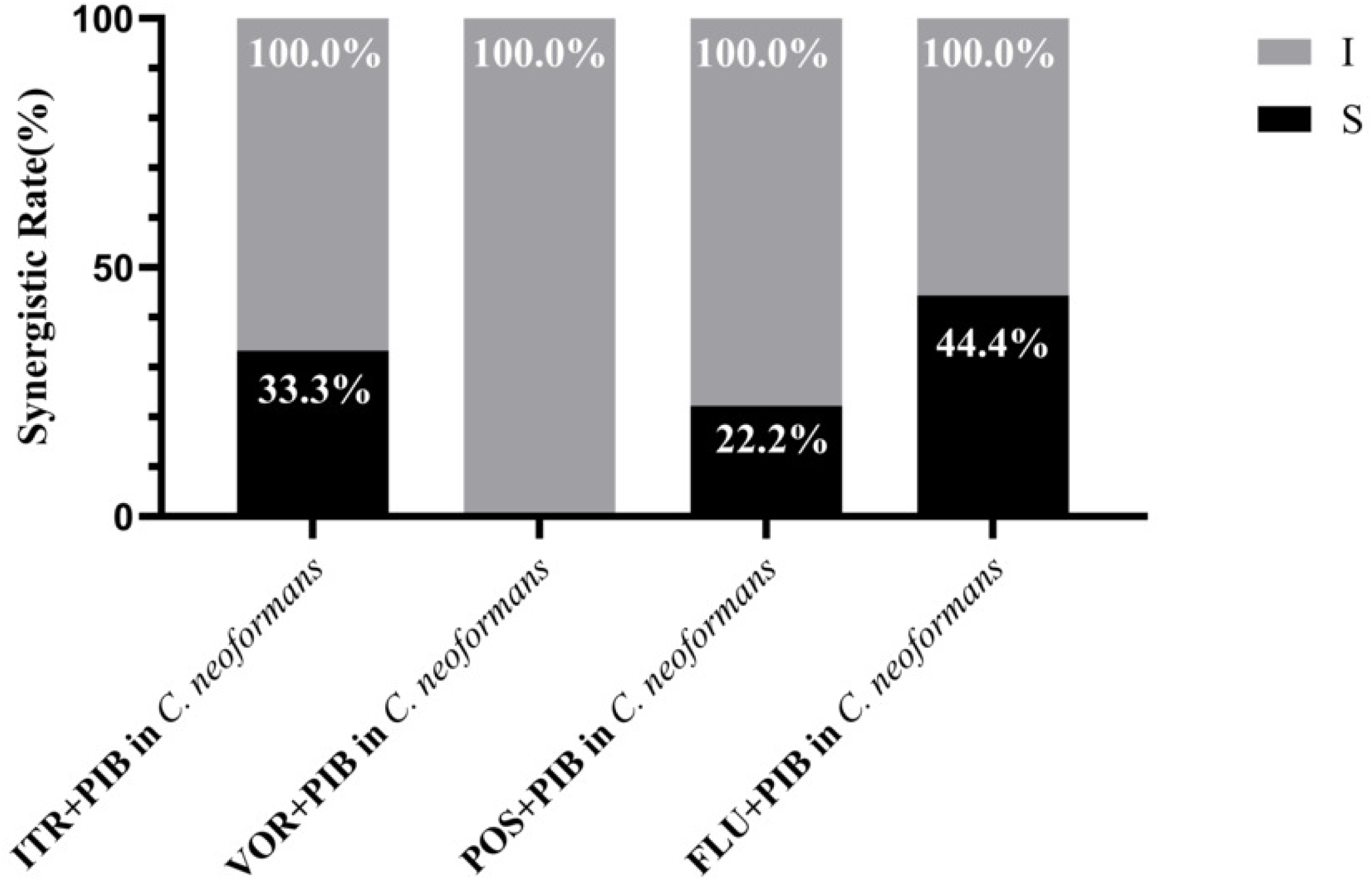
*lit vitro* drug sensitization results of PIB combined with azoles against *C.neoformans*. Summary of drug interaction for the combination of PIB and azoles. (A) Summary of interaction relationships for all drug combinations against *C.neoformans.* The fraction of in vitro interaction results of PIB combined with total result in *C.neoformans,* ITR, VOR, POS and FLU antifungal agents, respectively.S, synergy (FICI of < 0.5); I, no interaction (indifference)(0.5 < FICI < 4).

**Table 3.** *In vitro* drug sensitization results of PIB combined with azoles against *C. neoformans*.

| Strains | Alone |  |  |  |  | Combination |  |  |  |
| --- | --- | --- | --- | --- | --- | --- | --- | --- | --- |
|  | PIB | ITR | VOR | POS | FLU | PIB/ITR | PIB/VOR | PIB/POS | PIB/FLU |
| G10 | >16 | 0.25 | 0.125 | 0.25 | 0.5 | 1/0.125(I) | 0.125/0.12(I) | 0.25/0.125(I) | 1/0.25(I) |
| G11 | >16 | 0.25 | 0.125 | 0.25 | 1 | 0.5/0.12(I) | 0.125/0.12(I) | 2/0.063(S) | 0.25/1(I) |
| G12 | >16 | 0.5 | 0.125 | 0.25 | 0.25 | 2/0.125(I) | 0.5/0.063(I) | 2/0.125(I) | 0.5/0.063(S) |
| Z3 | >16 | 0.125 | 0.125 | 0.125 | 1 | 0.125/0.125(I) | 0.125/0.125(I) | 0.125/0.125(I) | 0.125/1(I) |
| Z4 | >16 | 0.5 | 0.125 | 0.25 | 1 | 0.25/0.25(I) | 0.125/0.125(I) | 0.5/0.125(I) | 4/0.25(S) |
| Z5 | >16 | 0.25 | 0.125 | 0.25 | 1 | 2/0.063(S) | 1/0.063(I) | 0.5/0.125(I) | 0.25/1(I) |
| Z6 | >16 | 0.25 | 0.125 | 0.25 | 0.25 | 0.25/0.125(I) | 0.125/0.125(I) | 0.5/0.063(I) | 4/0.063(S) |
| Z8 | >16 | 0.25 | 0.125 | 0.125 | 0.5 | 2/0.063(S) | 0.125/0.125(I) | 1/0.63(I) | 1/1(I) |
| Z9 | >16 | 0.25 | 0.125 | 0.25 | 1 | 1/0.125(I) | 0.125/0.125(I) | 4/0.063(S) | 4/0.25(S) |
| **Quality control** |  |  |  |  |  |  |  |  |  |
| ATCC22019 | >16 | 0.25 | 0.125 | 0.063 | 1 | 0.5/0.063(S) | 0.125/0.125(I) | 1/0.031(I) | 2/0.5(I) |

Note: ITR, itraconazole; VOR, voriconazole; POS, posaconazole; FLU; Fluconazole PIB, pibrentasvir; S, synergy (FICI $\leq 0.5$ ); I, indifference (no interaction, FICI from $>0.5$ to $\leq 4$ ). MICs were the concentrations that achieved 100% growth inhibition; FICI: fractional inhibitory concentration index.

### *In Vitro* Interactions of PIB in Combination with Azoles against *E. dermatitidis*

PIB alone showed no detectable antifungal activity against any of the 18 *E. dermatitidis* isolates, with MICs of > 16 μg/mL. The MIC ranges of ITR, VOR, and POS were 0.25-8 μg/mL, 0.125-8 μg/mL, and 0.125-8 μg/mL, respectively. Synergistic interactions were observed for PIB-ITR and PIB-POS against 10 (55.6%) and 11 (61.1%) isolates, respectively, whereas PIB-VOR exhibited synergy against only 2 isolates (11.1%). No antagonistic interactions were detected (Table 4). Among the tested combinations, PIB-POS showed the highest rate of synergy against *E. dermatitidis*, indicating a favorable *in vitro* interaction between PIB and POS (Figure 4).

**Figure 4.**
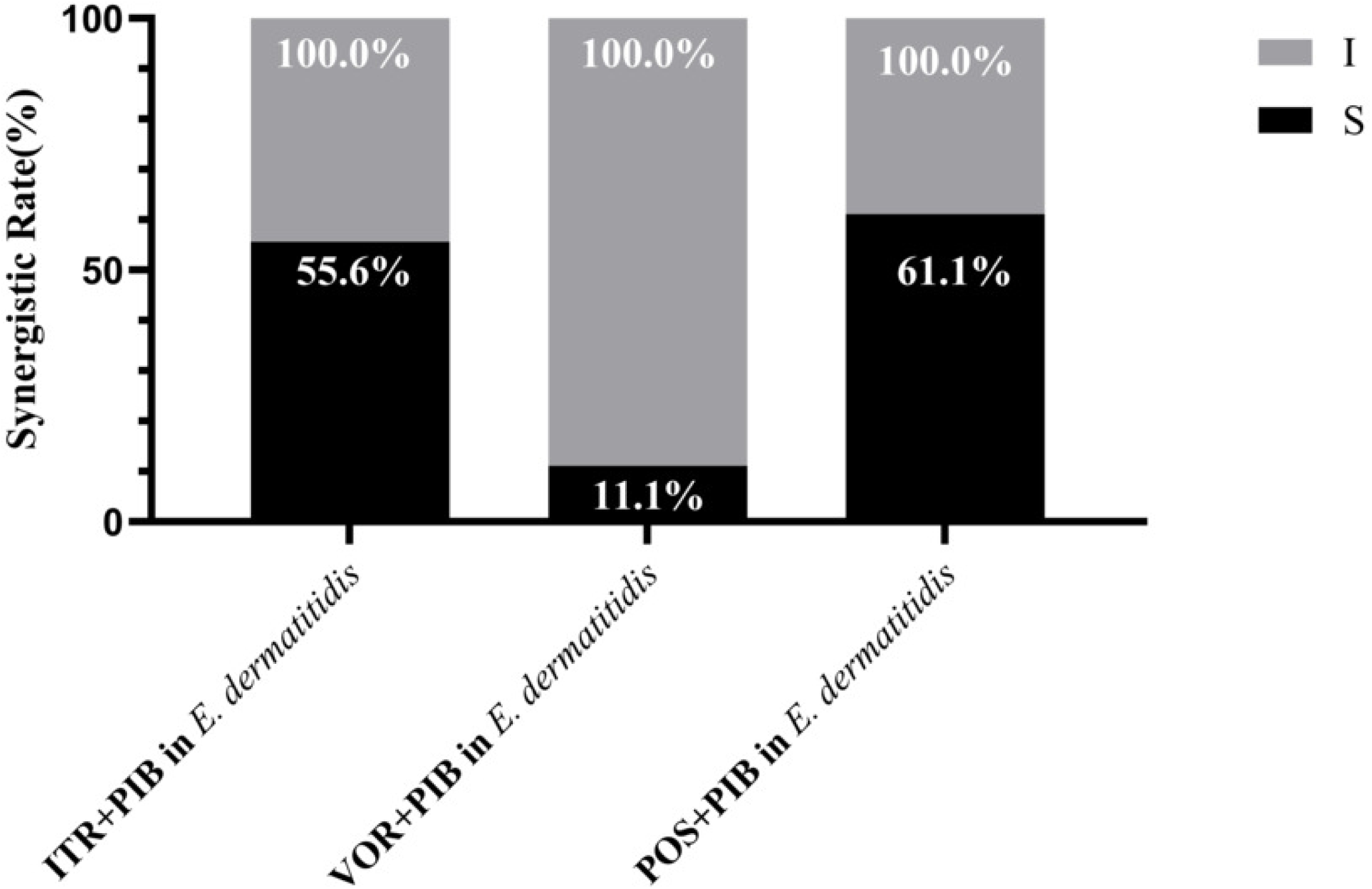
*lit vitro* drug sensitization results of PIB combined with azoles against *E.dermatitidis.* Summary of drug interaction for the combination of PIB and azoles. (A) Summary of interaction relationships for all drug combinations against *E.dermatitidis.* The fraction of in vitro interaction results of PIB combined with total result in E.dermatitidis, ITR, VOR, and POS antifungal agents, respectively.S, synergy (FICI of < 0.5); I, no interaction (indifference)(0.5 < FICI < 4).

**Table 4.** *In vitro* drug sensitization results of PIB combined with azoles against *E. dermatitidis*.

| Strains | Alone |  |  |  | Combination |  |  |
| --- | --- | --- | --- | --- | --- | --- | --- |
|  | PIB | ITR | VOR | POS | PIB/ITR | PIB/VOR | PIB/POS |
| 28 | >16 | 0.5 | 0.25 | 0.25 | 1/0.125(S) | 1/0.125(I) | 4/0.063(S) |
| 29 | >16 | 0.5 | 0.5 | 0.25 | 1/0.25(I) | 2/0.25(I) | 2/0.125(I) |
| 31 | >16 | 0.5 | 0.5 | 0.5 | 0.5/0.125(I) | 0.125/0.5(I) | 0.5/0.125(S) |
|  | PIB | ITR | VOR | POS | PIB/ITR | PIB/VOR | PIB/POS |
| D9g | >16 | 0.5 | 0.25 | 0.125 | 0.5/0.25(I) | 4/0.125(I) | 4/0.063(I) |
| D9h | >16 | 0.5 | 0.5 | 0.25 | 1/0.125(I) | 4/0.125(S) | 4/0.063(S) |
| D9i | >16 | 0.5 | 0.5 | 0.25 | 0.5/0.063(S) | 8/0.125(I) | 8/0.063(I) |
| D9j | >16 | 0.25 | 0.125 | 0.125 | 2/0.125(S) | 2/0.063(I) | 1/0.031(S) |
| D9k | >16 | 0.25 | 0.125 | 0.125 | 2/0.125(S) | 4/0.063(S) | 1/0.063(I) |
| 34 | >16 | 0.5 | 0.25 | 0.125 | 0.5/0.125(S) | 0.125/0.25(I) | 0.5/0.031(S) |
| 35 | >16 | 0.25 | 0.25 | 0.125 | 0.25/0.063(S) | 0.125/0.25(I) | 1/0.031(S) |
| 36 | >16 | 0.25 | 0.125 | 0.125 | 0.5/0.125(I) | 1/0.063(I) | 0.125/0.063(I) |
| 37 | >16 | 0.25 | 0.25 | 0.25 | 0.5/0.0625(S) | 0.125/0.25(I) | 0.5/0.031(S) |
| 38 | >16 | 0.25 | 0.25 | 0.5 | 0.125/0.25(I) | 0.125/0.25(I) | 0.25/0.125(S) |
| 39 | >16 | 0.25 | 0.25 | 0.125 | 1/0.063(S) | 0.125/0.25(I) | 1/0.03125(S) |
| 40 | >16 | 0.5 | 0.25 | 0.125 | 1/0.125(S) | 0.125/0.25(I) | 0.5/0.063(I) |
| 41 | >16 | 0.25 | 0.125 | 0.25 | 4/0.063(S) | 8/0.063(I) | 0.5/0.063(S) |
| 109140 | >16 | 0.25 | 0.25 | 0.125 | 0.125/0.25(I) | 0.5/0.125(I) | 2/0.031(S) |
| 109144 | >16 | 8 | 8 | 8 | 8/8(I) | 8/8(I) | 8/8(I) |
| **Quality control** |  |  |  |  |  |  |  |
| ATC204304 | >16 | 0.5 | 0.25 | 0.125 | 0.5/0.25(I) | 0.125/0.25(I) | 0.5/0.063(I) |
| ATCC22019 | >16 | 0.25 | 0.125 | 0.063 | 0.5/0.063(S) | 0.125/0.125(I) | 1/0.031(I) |

Note: ITR, itraconazole; VOR, voriconazole; POS, posaconazole; PIB, pibrentasvir; S, synergy (FICI $\leq 0.5$ ); I, indifference (no interaction, FICI from $>0.5$ to $\leq 4$ ). MICs were the concentrations that achieved 100% growth inhibition; FICI: fractional inhibitory concentration index.

**Fig. 5.**
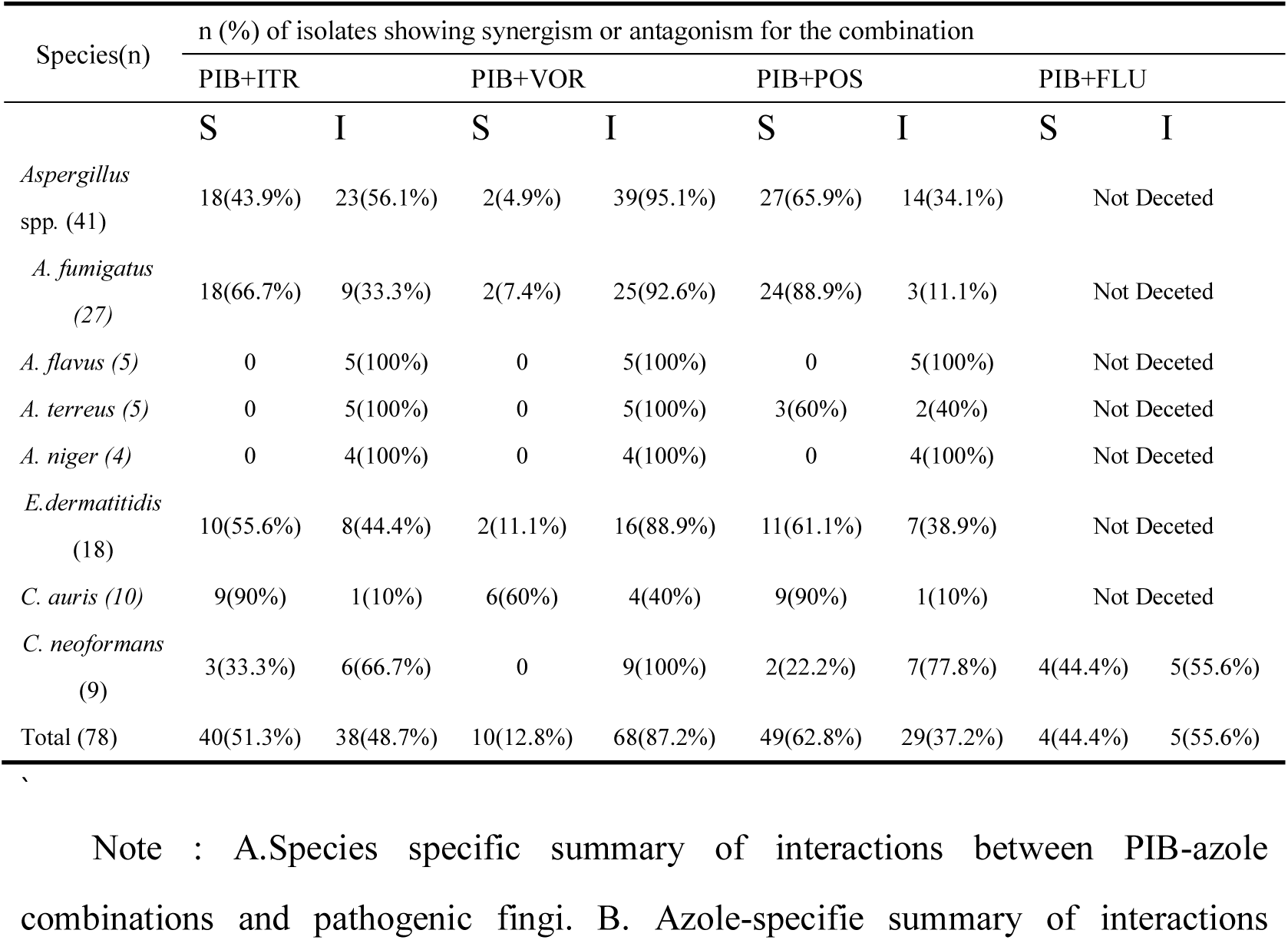
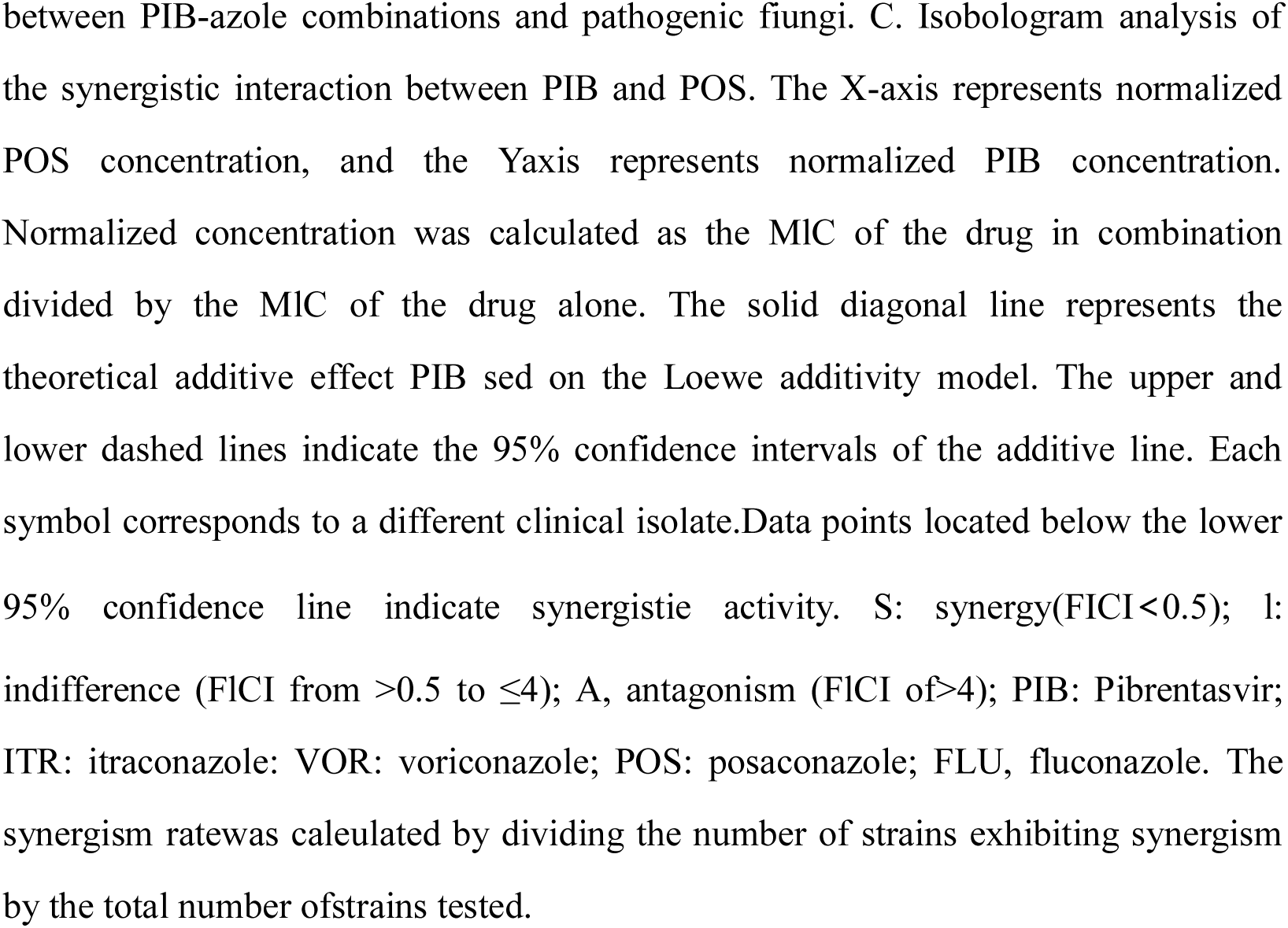
Summary of drug interaction for the combination of PIB and azole.

### Summary of the *In Vitro* Interactions between PIB and Azoles against Different Fungal Isolates

PIB in combination with antifungal agents exhibited substantial synergistic activity, although the frequency of synergy varied among fungal species and drug combinations. Combination therapy with PIB and POS produced the most widespread synergistic effects. Synergy was observed for 27 of 41 *Aspergillus* isolates (65.9%), 9 of 10 *C. auris* isolates (90.0%), 2 of 9 *C. neoformans* isolates (22.2%), and 11 of 18 *E. dermatitidis* isolates (61.1%). No antagonistic interactions were detected for the PIB-POS combination. PIB-ITR also demonstrated considerable synergistic activity against *Aspergillus* spp. (18/41, 43.9%), *C. auris* (9/10, 90.0%), *C. neoformans* (3/9, 33.3%), and *E. dermatitidis* (10/18, 55.6%), with no antagonism observed. The PIB-FLU combination exhibited synergistic activity against 4 of 9 *C. neoformans* isolates (44.4%). In contrast, PIB-VOR showed relatively limited synergistic activity, with synergy observed against *Aspergillus* spp. (2/41, 4.9%), *C. auris* (6/10, 60.0%), and *E. dermatitidis* (2/18, 11.1%). Nevertheless, no antagonistic interactions were detected for either the PIB-VOR or PIB-FLU combinations.

When all tested fungal isolates were analyzed collectively, the overall synergistic rates of PIB combined with ITR, VOR, and POS were 51.3% (40/78), 12.8% (10/78), and 62.8% (49/78), respectively. Because FLU was evaluated only against *C. neoformans*, the synergistic rate of PIB-FLU was 44.4% (4/9). Overall, PIB combined with POS or ITR exhibited broader synergistic activity than PIB combined with VOR. Among all combinations, PIB-POS showed the highest overall synergistic rate and demonstrated activity against *Aspergillus* spp., *C. auris*, *C. neoformans*, and *E. dermatitidis*, without detectable antagonism, indicating broad-spectrum synergistic potential (Figure 5). PIB-ITR also exhibited substantial synergistic activity across all four fungal groups, whereas the synergistic activity of PIB-VOR was comparatively limited.

**Figure 5.**
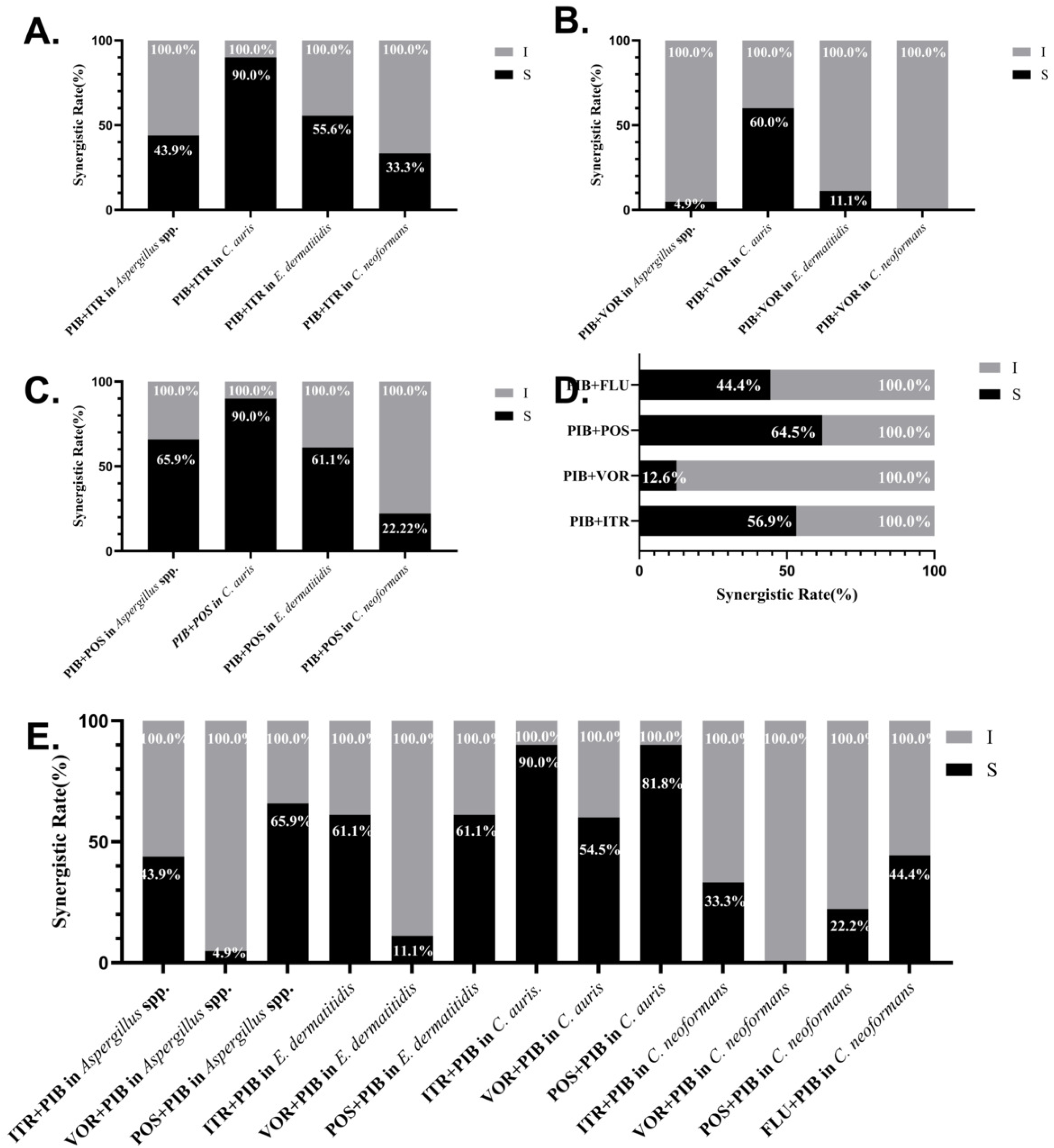
The Summary of the in vitro interactions between pibrentasvir (PIB) and azole antifungal agents. Note: (A-C) Distribution of synergistic and indifferent interactions of PIB combined with itraconazole (ITR), voriconazole (VOR), and posaconazole (POS), respectively, against *Aspergillus* spp., *C. auris, E. dermatitidis,* and C. *neoformans.* (D) Overall synergistic rates of PIB combined with ITR, VOR, POS, and FLU against all tested fungal isolates. (E) Summary of the interaction profiles of the indicated PIB-azoles combinations against each fungal species or group. Percentages within the bars indicate the proportions of isolates exhibiting the corresponding interaction. S, synergy (FICI < 0.5); I, indifference (0.5 < FICI <4).

## Discussion

IFIs are systemic infections caused by yeasts or molds that invade and establish infection in deep tissues. Unlike superficial fungal infections, IFIs are life-threatening diseases associated with substantial morbidity and mortality. The most common clinically relevant fungal pathogens belong to the genera *Candida*, *Aspergillus*, and *Cryptococcus*. In recent years, the incidence of infections caused by *E. dermatitidis* has shown a progressively increasing trend^19^. It is estimated that approximately 1.9 million people develop acute IFIs annually, while nearly 3 million people worldwide are affected by chronic severe fungal infections. Many of these infections are life-threatening and are estimated to cause more than 1.6 million deaths each year^20^. Meanwhile, the increasing prevalence of antifungal resistance and the continued decline in fungal susceptibility have substantially complicated the treatment of fungal infections. Combination therapy has emerged as a valuable strategy for preventing resistance and restoring antifungal efficacy, and has been successfully applied to the treatment of a variety of fungal infections, particularly those caused by multidrug-resistant fungal pathogens. PIB is a pan-genotypic inhibitor targeting the HCV NS5A protein. It has been reported to exhibit *in vitro* activity against HCV genotypes 1-6 and to retain activity against multiple common NS5A resistance-associated substitutions. To our knowledge, this study is the first to systematically demonstrate that PIB exerts significant synergistic antifungal effects when combined with azole agents.

Several studies have demonstrated that antiviral agents have considerable potential for antifungal drug development and represent promising candidates for combination therapy against fungal infections. *In vitro* studies have shown that various human immunodeficiency virus protease inhibitors (HIV-PIs), including atazanavir (ATV), saquinavir (SQV), lopinavir (LPV), and ritonavir (RTV), exhibit significant synergistic activity when combined with antifungal agents against invasive fungal infections, including candidiasis and cryptococcosis. Moreover, these combination regimens have generally demonstrated favorable safety profiles^21^. The chemosensitizing activity of these agents was superior to that of several previously reported azole sensitizers, including sulfamethoxazole, clorgyline, and cyclosporine^22–24^. The observed synergistic effect may be associated with the inhibition of hyphal growth and biofilm formation^25,26^.

Similarly, the HCV NS5A inhibitors DCV and VEL, which share a mechanism of action similar to that of PIB, also exhibit potential as antifungal sensitizers. Both agents form consistent synergistic antifungal combinations with AmB, showing activity not only against multiple pathogenic Mucorales species but also against other invasive fungal pathogens, including *A. fumigatus*, *C. neoformans*, and *C. auris*. This synergistic effect may be associated with the disruption of cellular cholesterol trafficking^11^. Given the mechanistic similarity of PIB to DCV and VEL, its antifungal activity may be mediated through comparable pathways. In addition, pharmacokinetic studies have shown that PIB can weakly inhibit the hepatic uptake transporters OATP1B1/3, the metabolic enzyme CYP3A, and the efflux transporters P-glycoprotein (P-gp) and breast cancer resistance protein (BCRP). Azole antifungal agents are also substrates of OATP1B1/3, CYP3A, and P-gp/BCRP. Therefore, we speculate that the coadministration of PIB may reduce the systemic clearance of azoles, thereby increasing azole exposure and potentially enhancing their antifungal activity^27^.

Although PIB alone exhibited no detectable antifungal activity, its combination with ITR, VOR, POS, or FLU produced synergistic effects against *Aspergillus* spp., *C. auris*, *E. dermatitidis*, and *C. neoformans*. The synergistic rates observed for PIB combined with ITR or POS were generally higher than those observed with VOR or FLU, which may be related to the long side chains and high hydrophobicity of POS and ITR^28^. Notably, the PIB-POS combination exhibited the highest overall synergistic activity, with synergy observed against *Aspergillus* spp. (65.9%), *C. aruis* (90.0%), *E. dermatitidis* (61.1%), and *C. neoformans* (22.2%). No antagonistic interactions were detected, suggesting that this combination has broad-spectrum synergistic potential. ITR and POS possess complex molecular side chains that enable high-affinity binding to fungal lanosterol CYP51, thereby potently inhibiting the ergosterol biosynthetic pathway. Their antifungal activities are generally greater than those of FLU and VOR. As a structurally optimized derivative of ITR with an extended side chain, POS contains a longer hydrophobic piperazinyl-aryl moiety, which expands its interactions within the CYP51-binding pocket and may consequently enhance its overall antifungal potency and broaden its spectrum of activity^28–30^.

This study has several limitations. First, the investigation was restricted to *in vitro* experiments, and the findings may not fully reflect the pharmacological behavior or therapeutic potential of these compounds in vivo. Future studies should therefore evaluate the efficacy of these combinations in appropriate animal models and further elucidate their underlying mechanisms, thereby providing a stronger theoretical basis for the rational and precise use of PIB in combination with antifungal agents. Overall, PIB exhibited marked synergistic activity when combined with antifungal drugs, highlighting the potential of antiviral drug repurposing as a strategic approach to addressing the urgent unmet therapeutic needs associated with invasive fungal infections. These findings also provide a valuable starting point for further investigation. PIB and related compounds may serve as promising scaffolds for medicinal chemistry optimization or for the identification of novel antifungal mechanisms, which may ultimately facilitate the development of safer and more effective therapeutic strategies for these difficult-to-treat infections.

## 5. Declarations

### Ethics approval

Not applicable.

### Consent for publication

Not applicable.

### Availability of data and materials

Not applicable.

### Author contributions

All authors contributed to the research in this report. Individual contributions are as follows: Conceptualization, YS and HZ; methodology, JH; software, XYT and HMY; validation, JH; formal analysis, LG; investigation, HMY; resources, YS; data curation, SJL; writing original draft preparation, HZ; writing review and editing, YS and HZ; visualization, SJL and JH; supervision, XLZ; project administration, YS; funding acquisition, YS and HZ. All authors have read and agreed to the published version of the manuscript.

## Acknowledgments

We thank everyone who contributed to the success of this research, including colleagues, institutions, and funding bodies.

## References

1. Brown GD, Denning DW, Gow NAR et al. Hidden Killers: Human Fungal Infections. Sci Transl Med. 2012;4(165):165rv13–165rv13

2. Latgé J. Aspergillus fumigatus and Aspergillosis. Clin Microbiol Rev. 1999;12(2):310–350

3. Firacative C. Invasive fungal disease in humans: are we aware of the real impact? Memórias do Instituto Oswaldo Cruz. 2020;115:e200430

4. Denning DW. Global incidence and mortality of severe fungal disease. The Lancet Infectious Diseases. 2024;24(7):e428–e438

5. Alanio A, Snell HM, Cordier C et al. First Patient-to-Patient Intrahospital Transmission of Clade I Candida auris in France Revealed after a Two-Month Incubation Period. Microbiol Spectr. 2022;10(5):e0183322

6. Murthy JMK, Sundaram C. Chapter 95 – Fungal infections of the central nervous system. 2014;121:1383–1401

7. Suchodolski J, Parol M, Pawlak K, Piecuch A, Ogórek R. Clinical and Molecular Advances on the Black Yeast Exophiala dermatitidis. International Journal of Molecular Sciences. 2025;26(14):6804

8. Binjubair FA, Parker JE, Warrilow AG et al. Small-Molecule Inhibitors Targeting Sterol 14α-Demethylase (CYP51): Synthesis, Molecular Modelling and Evaluation Against Candida albicans. ChemMedChem. 2020;15(14):1294–1309

9. Fisher MC, Alastruey-Izquierdo A, Berman J et al. Tackling the emerging threat of antifungal resistance to human health. Nature reviews. Microbiology. 2022;20(9):557–571

10. Johnson MD, MacDougall C, Ostrosky-Zeichner L, Perfect JR, Rex JH. Combination antifungal therapy. Antimicrob Agents Chemother. 2004;48(3):693–715

11. Khan AA, AlKashef NM, Seleem MN. Repurposing antiviral agents against mucormycosis. PLoS One. 2026;21(2):e0342559

12. Glass NL, Donaldson GC. Development of primer sets designed for use with the PCR to amplify conserved genes from filamentous ascomycetes. Appl Environ Microbiol. 1995;61(4):1323–1330

13. Institute CALS. Reference Method for Broth Dilution Antifungal Susceptibility Testing of Yeasts; Approved Standard—Third Edition. 2008;(1-56238-666-2)

14. Institute CALS. Reference Method for Broth Dilution Antifungal Susceptibility Testing of Filamentous Fungi; Approved Standard—Second Edition. 2008;(1-56238-668-9)

15. Gao L, Xia X, Gong X, Zhang H, Sun Y. In vitro interactions of proton pump inhibitors and azoles against pathogenic fungi. Front Cell Infect Microbiol. 2024;14:1296151

16. Jia G, Hu J, Tan L et al. In Vitro and In Vivo Evaluation of Synergistic Effects of Everolimus in Combination with Antifungal Agents on Exophiala dermatitidis. Microbiol Spectr. 2023;11(3):e05302–22

17. Tobudic S, Kratzer C, Lassnigg A, Graninger W, Presterl E. In vitro activity of antifungal combinations against Candida albicans biofilms. The Journal of antimicrobial chemotherapy. 2010;65(2):271–274

18. Fothergill AW. Antifungal Susceptibility Testing: Clinical Laboratory and Standards Institute (CLSI) Methods. 2012:65–74

19. Fang W, Wu J, Cheng M et al. Diagnosis of invasive fungal infections: challenges and recent developments. J Biomed Sci. 2023;30(1):42

20. Bongomin F, Gago S, Oladele RO, Denning DW. Global and Multi-National Prevalence of Fungal Diseases—Estimate Precision. J Fungi (Basel). 2017;3(4):57

21. Elgammal Y, Salama EA, Seleem MN. HIV protease inhibitors restore amphotericin B activity against Candida. PLoS One. 2025;20(5):e0324080

22. Eldesouky HE, Li X, Abutaleb NS, Mohammad H, Seleem MN. Synergistic interactions of sulfamethoxazole and azole antifungal drugs against emerging multidrug-resistant Candida auris. Int J Antimicrob Agents. 2018;52(6):754–761

23. Marchetti O, Moreillon P, Glauser MP, Bille J, Sanglard D. Potent Synergism of the Combination of Fluconazole and Cyclosporine in Candida albicans. Antimicrob Agents Chemother. 2000;44(9):2373–2381

24. Holmes AR, Keniya MV, Ivnitski-Steele I et al. The Monoamine Oxidase A Inhibitor Clorgyline Is a Broad-Spectrum Inhibitor of Fungal ABC and MFS Transporter Efflux Pump Activities Which Reverses the Azole Resistance of Candida albicans and Candida glabrata Clinical Isolates. Antimicrob Agents Chemother. 2012;56(3):1508–1515

25. Alkashef NM, Seleem MN. Novel combinatorial approach: Harnessing HIV protease inhibitors to enhance amphotericin B’s antifungal efficacy in cryptococcosis. PLoS One. 2024;19(8):e0308216

26. Elgammal Y, Salama EA, Seleem MN. Saquinavir potentiates itraconazole’s antifungal activity against multidrug-resistant Candida auris in vitro andin vivo. Med Mycol. 2023;61(9):myad081

27. Kosloski MP, Oberoi R, Wang S et al. Drug-Drug Interactions of Glecaprevir and Pibrentasvir Coadministered With Human Immunodeficiency Virus Antiretrovirals. The Journal of infectious diseases. 2020;221(2):223–231

28. Shi N, Zheng Q, Zhang H. Molecular Dynamics Investigations of Binding Mechanism for Triazoles Inhibitors to CYP51. Front Mol Biosci. 2020;Volume 7 – 2020

29. Gupta AK, Tomas E. New antifungal agents. Dermatol Clin. 2003;21(3):565–576

30. Hof H. A new, broad-spectrum azole antifungal: posaconazole--mechanisms of action and resistance, spectrum of activity. Mycoses. 2006;49 Suppl 1:2–6

